# Collagen Shapes Fingertip Surface Strains during Normal Loading

**DOI:** 10.64898/2026.02.08.704563

**Authors:** Guillaume H. C. Duprez, Donatien Doumont, Philippe Lefèvre, Benoit P. Delhaye, Laurent Delannay

## Abstract

When making contact, fingertip mechanoreceptors respond to the skin deformation, and provide essential information for tactile perception and object manipulation. Since subsurface measurements remain challenging, strains close to the receptors are commonly estimated using numerical models. Here, we present a biomechanical finite element model simulating fingertip normal loading against a flat plate. Several model variants are designed to isolate the role of tissue heterogeneity and collagen-induced anisotropy. Their predictions are compared to experimental data of fingertip surface strains obtained with 3-D stereo imaging. By varying the stiffness contrast and fiber orientation, we demonstrate that incorporating collagen anisotropy is required to reproduce strain localization at the contact edge while maintaining realistic global shape changes. In particular, fibers aligned parallel to the skin surface induce local skin thickening and a pronounced radial expansion beneath the contact edge, affecting mechanoreceptors. This observation suggests a collagen-mediated contribution to the deep transmission of mechanical stimuli. These results highlight collagen architecture as a key determinant of fingertip mechanics and underscore its importance for accurate modeling of tactile interactions.

## 1. Introduction

When the fingertip makes contact with a surface, the skin deformation is a mechanical stimulus that activates cutaneous mechanoreceptors, giving rise to tactile perception (Johansson and Flanagan, 2009). Recently, optical coherence tomography (OCT) was used to image beneath the fingertip surface (Corniani et al., 2025). However, the speckle noise inherent to OCT imaging made it difficult to obtain reliable strain measurements at the depth of the mechanoreceptors. The present study aims to develop a biomechanical model of the fingertip deformation assessed based on experimental measurements of surface strains. Progress in imaging techniques now makes it possible to probe deformations over the entire external surface of the fingertip during normal loading (Doumont et al., 2025). Earlier studies primarily focused on the fingerpad (i.e., the part in contact), which was imaged under normal (Willemet et al., 2021), tangential (Bochereau et al., 2017; Delhaye et al., 2014), and torsional (du Bois de Dunilac et al., 2023) loading conditions, as well as during active object manipulation (Delhaye et al., 2021b). With the help of a physically-sound numerical model, the measured surface strains could be directly linked to the activity of underlying mechanoreceptors (Delhaye et al., 2021a).

Early efforts to model fingertip biomechanics relied on analytical approaches, primarily addressing simplified conditions such as line loads. As a first approximation, the surface deflection of the fingertip was roughly approached by treating the fingertip as an incompressible, homogeneous, semi-infinite half-space obeying Hooke’s law (Phillips and Johnson, 1981). A more refined approach, the *waterbed* model, represented the fingertip as a two-dimensional (2D) system consisting of a thin, flexible membrane (the skin) overlaying an incompressible fluid (subcutaneous tissue) (Srinivasan, 1989). The latter provided realistic predictions of surface deflection at the position of the loading, but its accuracy diminished significantly at other locations. Other studies have focused on the fingertip response when pressed against a flat plate. An axisymmetric ellipsoidal, inflated membrane was shown to effectively replicate load-displacement behavior, and contact area evolution (Serina et al., 1998). This highlighted the importance of geometry, material heterogeneity, and skin’s natural tension (Annaidh and Destrade, 2019) in capturing fingertip biomechanics.

More recent models of the fingertip’s biomechanics relied on the finite element (FE) method. The latter allowed the study of more complex loadings while accounting for the heterogeneity of the stress and strain fields (see for review Cei et al. (2025)). Simple FE representations of the fingertip considered homogeneous, incompressible, linear-elastic cross-sections of the pulp under the assumption of plane-strains (Srinivasan and Dandekar, 1996). These simplified models were shown to be less accurate than the waterbed model, further emphasizing the influence of the contrasted response of the skin relative to the subcutaneous tissues. Improvements were later achieved by introducing a bilayer fingertip structure with isotropic, non-linear (hyperelastic) material behavior (Wu et al., 2004). The interaction between the fingertip and a flat plate was investigated using both 2D (Serhat et al., 2022) or three-dimensional (3D) (Harih and Tada, 2015) isotropic models. These models were evaluated based on the accuracy of their predicted force-indentation curves (Wu et al., 2006) and the growth of the contact area (Hokari and Pramudita, 2023). Some studies also considered the distribution of contact pressure (D’Angelo et al., 2017). In a previous work (Duprez et al., 2024), some of the present authors relied on improved constitutive equations for the modeling of biological tissues (see for review Holzapfel and Ogden (2025)). They showed that considering the anisotropy of the subcutaneous network of collagen fibers (Hauck et al., 2004) allows a better reproduction of the contact area and contact pressure. The fingertip was then modeled as a homogeneous hyperelastic and anisotropic material.

It remains unclear how the heterogeneity of fingertip tissues, combined with the anisotropy arising from the collagen fiber network, influence surface strain patterns. The dermis (i.e., the thickest layer of the skin) is mainly composed of collagen fibers (Daly, 1982) running parallel to the skin surface (Munisso et al., 2023; Silver, 2022). While an idealized representation of the dermal fiber network was shown to faithfully reproduce the biaxial response of skin under tensile load (Gibson et al., 1965), dermal collagen has never been considered when modeling fingertip interactions. These fibers have, however, been shown to be the main source of tensile strength in the skin (Haut, 2002).

In this work, we aim to characterize the influence of several intrinsic biomechanical properties of the fingertip skin on its response to normal loading. More precisely, we assess the influence of heterogeneity (i.e., layered structure) and presence of collagen inducing both anisotropy and traction–contraction asymmetry. Comparing simulations with experimental data guides the refinement of the model towards a more “physiological” fingertip representation which combines these individual biomechanical characteristics.

## 2. Experimental data

In the experiment shown in Figure 1 (see for details Doumont et al. (2025)), the index fingertip of nine subjects was loaded against a flat glass plate from 0 N to 5 N at constant speed ∼5 mm/s (see Figure 1 (a)). Ink-sprayed features were tracked via 3D digital image correlation (3D-DIC), allowing a point cloud reconstruction of the volar surface of the fingertip under load (see Figure 1 (b)), and computation of the surface Green-Lagrange strain tensor **E**^GL^. The two principal strains E^GL^ and E^GL^ were found to align closely with the local meridional and circumferential directions, respectively (see Figure 1 (c)). As the distal fingertip region was close to half-spherical (see Figure 1 (b)), the value of **E**^GL^ was averaged circumferentially to observe its evolution along the meridional direction, hence assuming an axisymmetric behavior. The meridional 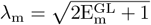and circumferential 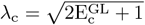 were averaged across subjects.

**Figure 1.**
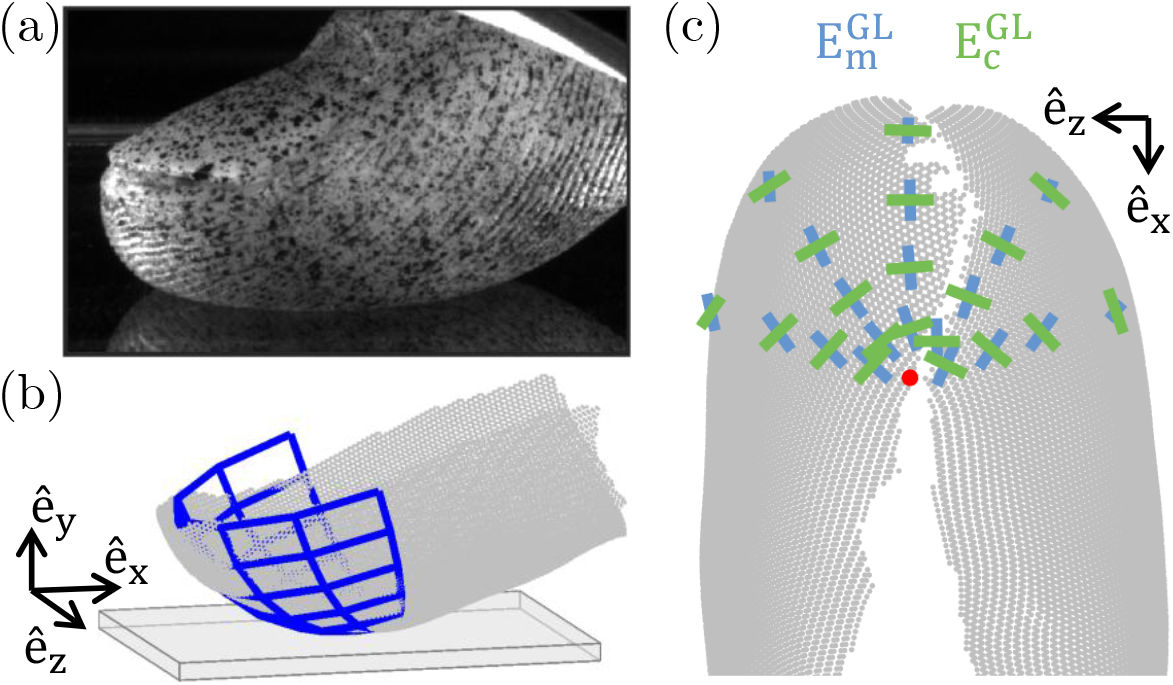
Description of the experiment made by Doumont et al. (2025). (a) Imaging of the fingertip surface covered with sprayed-ink during normal loading; (b) Spherical fit to the fingertip point-cloud reconstruction before loading; (c) Local principal directions of the Green-Lagrange strains.

The experimental data is revisited here to analyse the pulp deformation more globally. An axes-aligned spheroid is fitted to the distal region of each subject’s 3D fingertip point cloud as described in Figure 2. The frame-to-frame variation of the spheroid radius perpendicular to the loading direction is used as an approximation of the fingertip lateral expansion. The region of contact with the rigid plate is identified as an ellipse in the x–z plane (Figure 2 (a)). Coplanar points within the ellipse are removed to prevent bias in the spheroid fit. The x-coordinate of the ellipse center (see black dot in Figure 2 (a)) is used as a threshold to isolate the fingertip’s most distal region in each frame.

**Figure 2.**
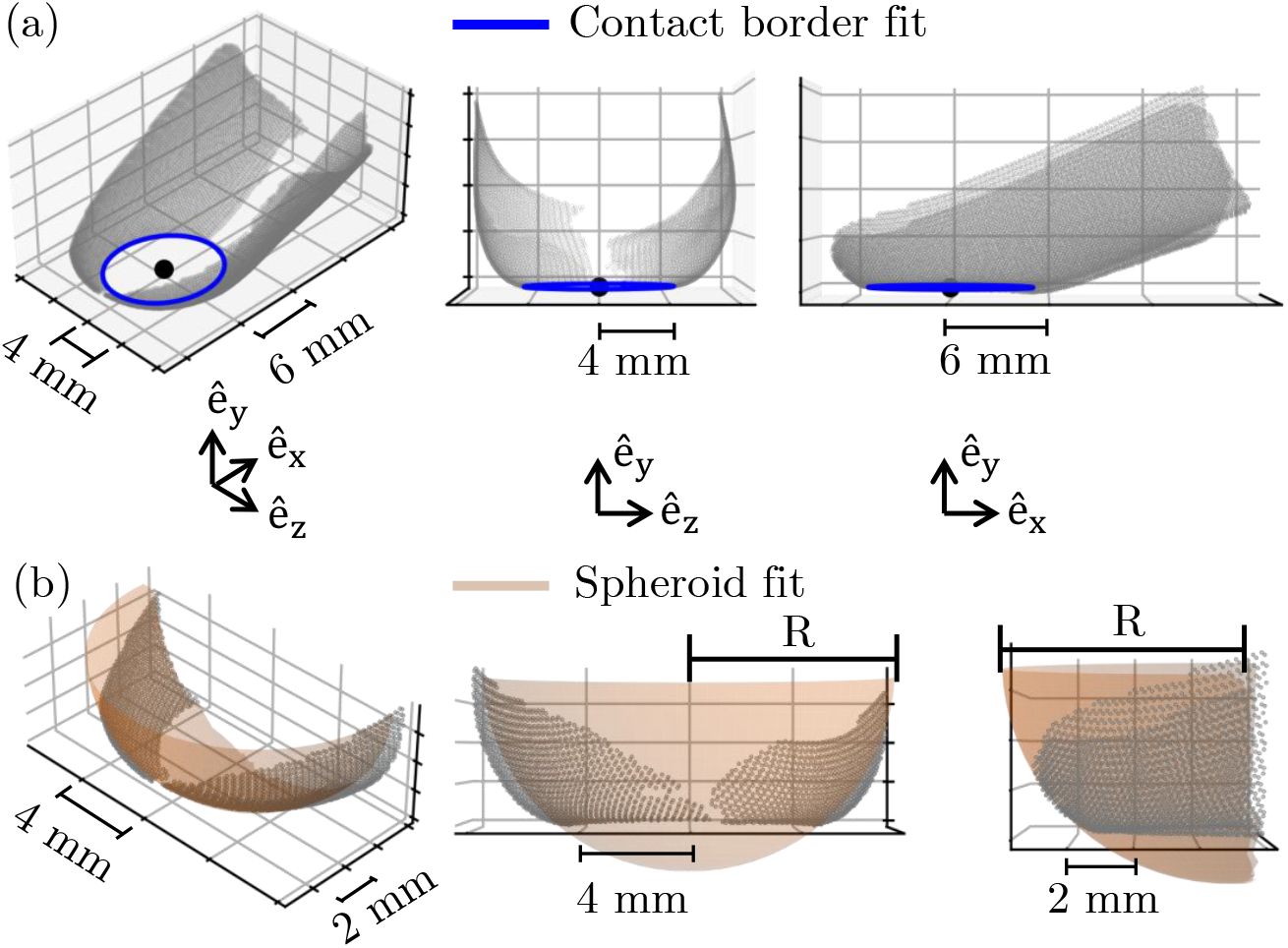
Description of the estimation of the evolution of the shape of the fingertip pulp during loading using 3D point-cloud reconstruction of subjects’ fingertips. (a) Fit of an ellipse to the border of the contact region with highlighted ellipse center; (b) Fit of a spheroid to the distal fingertip region.

The fit shown in Figure 2 (b) is obtained by minimizing the cost function J, defined as

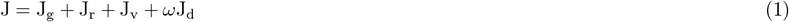

where J_g_ is the mean squared distance of the measured points to the spheroid, while J_r_, J_v_ and J_d_ act as regularizers. Specifically, J_r_ penalizes time fluctuations in the spheroid radii. It is calculated as the sum of squared relative changes between consecutive frames. Similarly, J_v_ constrains variations in the spheroid volume above the contact plane by measuring the squared relative deviation from its initial value. The final regularization term, J_c_, penalizes motion of the spheroid center transverse to the loading direction. It is defined as the sum of squared relative deviations of the x and z coordinates from their initial values (see axes in Figure 2). The dimensionless scalar weight *ω* = 4 is introduced to limit lateral displacement and is selected empirically. To reduce sensitivity to initialization while still exploring the search space effectively, the bounded problem is first solved with a particle swarm optimization (PSO) scheme implemented via the Python `pyswarms` library. The best PSO solution then seeds a second-stage refinement using the L-BFGS-B algorithm from the Python `scipy` library.

Figure 3 (a) shows that the fit captures an increase in the spheroid radius during loading. As expected, its displacement occurs primarily along the loading direction. ê_y_ (see Figure 3 (b)). The resulting load-displacement relationship is consistent with those reported in previous experiments (Dzidek et al., 2017). A similar relationship is obtained from the relative motion between points identified as being in contact (i.e., approximately coplanar) and points near the nail, which were minimally affected by the applied load. Figure 3 (c) highlights that the spheroid radius and portion of volume below the contact plane increase. However, its volume above the contact plane remains stable. These observations suggest that the fingertip undergoes a slight lateral expansion during normal loading while preserving a constant volume.

**Figure 3.**
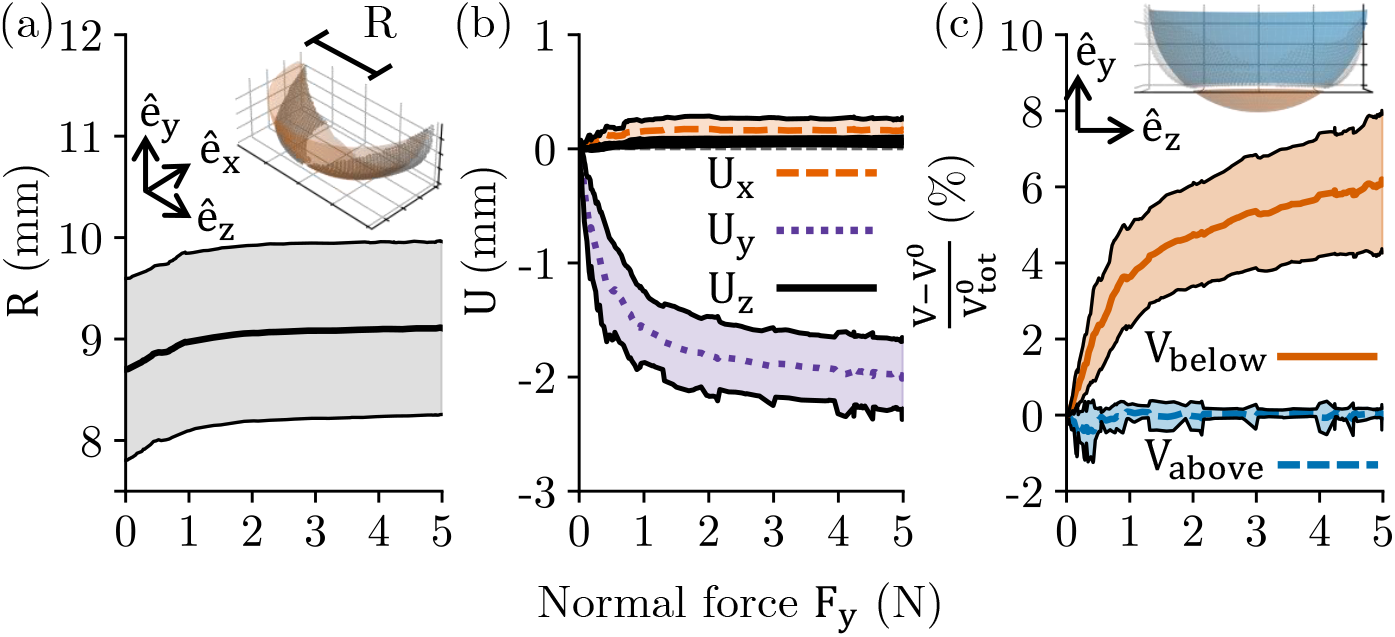
Results of the spheroid fit to the experimental point-cloud data of fingertip normal loading averaged over the nine subjects (with shaded standard deviation); (a) Evolution of the fitted spheroid radius in the x-z plane; (b) Displacement of the fitted spheroid center along the x-, y- and z-axes; (c) Relative changes in the fitted spheroid volume above and below the contact plane.

## 3. Description of the numerical model

The FE model proposed here relies on a simplified fingertip geometry. As shown in Figure 4 (a), the fingertip is considered semi-spherical since this reasonably approximates its most distal region (see Figure 1 (b)). The initial radius R is set to 7.5 mm which corresponds to half of the average height of male index fingertip (Serina et al., 1997). Because the experimental trends are described in terms of circumferential averages, the 3D problem is solved using a 2D axisymmetric formulation, taking ê_y_ as the revolution axis. Only the lower half of the fingertip is considered to deform. The vertical displacement of the top surface is impeded, but this surface is free to expand horizontally. It is thus assumed that there is no constraining effect of the nail at this depth across the finger.

**Figure 4.**
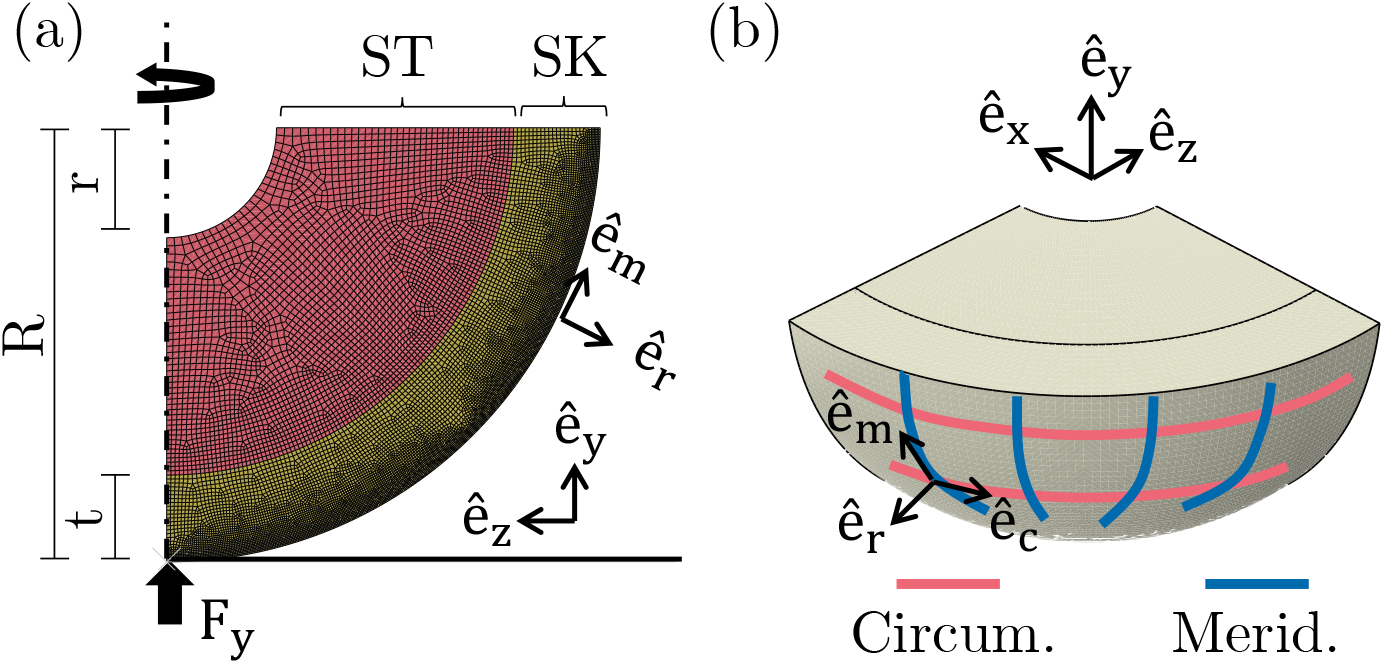
Description of the 2D axisymmetric FE model of fingertip normal loading. (a) Spherical fingertip made of two materials representing the subcutaneous tissues (ST) and skin (SK) in contact with a rigid flat plate; (b) Local meridional and circumferential directions along the surface of the model.

At the start of the simulation, the fingertip is positioned on a horizontal, rigid plate with zero initial contact pressure. A vertical force F_y_ is then applied to the plate to initiate its displacement along the positive ê_y_ direction. To reduce the number of model parameters, an infinite friction coefficient is assumed between the plate and the fingertip. This prevents relative motion between the two surfaces once contact is established. This behavior does not conform with the experimental observations, as will be discussed in the following Sections.

The fingertip is assumed to be made up of subcutaneous tissues (see ST in Figure 4 (a)) encapsulated in a thick skin layer (see SK in Figure 4 (a)). Both materials are considered incompressible. Although skin is typically divided into dermis and epidermis, it is modeled here as a single material with a material response different from the subcutaneous tissues. This is a reasonable assumption as the skin’s mechanical behavior is dominated by the dermis (Wahlsten et al., 2023). The thickness of the outer layer is fixed at 20% of the fingertip radius, which is a realistic approximation of the total skin thickness (Serhat et al., 2022). The bone is considered infinitely stiff. Accordingly, a spherical hole of radius r is made at the center of the spherical fingertip and the motion of the inner surface is impeded. The hole radius is fixed at 1.9 mm to match half of the fingertip bone height (Darowish et al., 2015).

Inside the spherical fingertip, the soft tissue core and skin outer layer are both assumed to behave as hyperelastic materials reinforced by collagen fibers along N directions. We rely on the incompressible form of the Holzapfel-Gasser-Ogden (HGO) anisotropic strain energy density function W_HGO_ (Gasser et al., 2006), which is given by

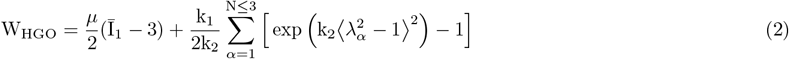

where the first term represents the elastic energy stored during the deformation of the isotropic background matrix, while the second term corresponds to the elastic energy stored in the collagen fibers. In the first term, the dimensionless scalar variable 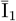 is the first invariant of the isochoric part of the right Cauchy-Green strain tensor 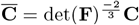 with **C** = **F**^**T**^**F** and the scalar variable *µ* (MPa) is the initial shear modulus of the material. In the second term of W_HGO_, the scalar k_1_ (MPa) represents the stiffness of collagen fibers, while the dimensionless scalar variable k_2_ characterizes their stiffening rate under stretch. The scalar 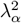 measures the square of the stretch (or contraction) of collagen fibers aligned with the unit vector a_*α*_ as 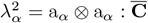. Finally, the Macaulay brackets condition ⟨ · ⟩ ensures that only fiber elongation contributes to the elastic energy, thereby inducing the traction–compression asymmetry (Holzapfel and Ogden, 2015).

Quasi-static simulations are carried out using Abaqus/Standard 2021 (*Simulia, Dassault Systemes*, France) FE solver with generalized axisymmetric elements. When dealing with incompressible materials such as soft biological tissues, only the deviatoric stresses and strains can be derived with the standard FE displacement formulation. Hence, the hybrid element formulation is used (CGAX4H), which can compute the pressure by enforcing volume preservation. The 2D axisymmetric quadrilateral mesh is generated using GMSH (Geuzaine and Remacle, 2009), allowing selective refinement near the contact region with the plate. The circumferential *λ*_c_, meridional *λ*_m_ and radial *λ*_r_ principal stretches are derived from the Lagrange strains.

Several variants of the model are adopted to investigate how different biomechanical properties influence the evolution of fingertip surface strains during normal loading (see Figure 5).

**Figure 5.**
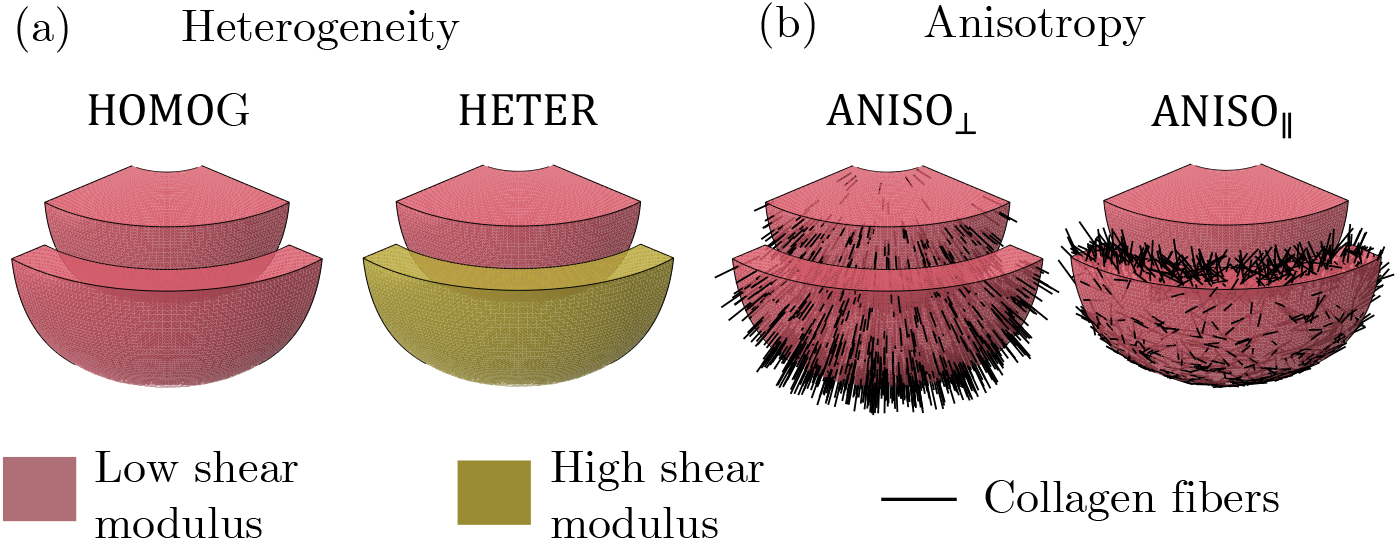
Variants of the model. (a) Isotropic variants; (b) Variants of the model reinforced by collagen fibers oriented perpendicular (or parallel to the outer surface).

- The effect of heterogeneity is evaluated by comparing the predictions obtained while relying on different ratios of the shear moduli of the skin layer *µ*^SK^ relative to the subcutaneous tissues *µ*^ST^. Both materials are considered isotropic 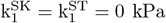. In the first variant, labeled HOMOG in Figure 5 (a), identical material properties are assigned to the skin and subcutaneous tissue: *µ*^SK^*/µ*^ST^ = 1. In the second variant, labeled HETER, a stiff skin encapsulates the more compliant subcutaneous tissue. Relying on Maeno et al. (1998), the ratio of shear moduli should be set to *µ*^SK^*/µ*^ST^ = 2.5 (HETER_2.5_). Since the range of reported dermal stiffness is large (Wahlsten et al., 2023), a greater shear moduli ratio is also tested *µ*^SK^*/µ*^ST^ = 10 (HETER_10_).
- The impact of anisotropy induced by collagen fibers is evaluated with additional model variants (see Figure 5 (b)). In both of them, the background matrix has the same shear modulus in the inner and outer layers (i.e., *µ*^SK^*/µ*^ST^ = 1). However, collagen fibers are introduced selectively in the two layers. The variant labeled ANISO_⊥_ investigates the influence of the anisotropy induced by collagen fibers radiating from the bone to the basement membrane of the epidermis (Hauck et al., 2004). Since skin is assumed as a single layer and the epidermis is a relatively thin outer skin layer, these fibers are considered in both inner and outer layer of the variant. To do so, a single radial collagen direction is prescribed at each integration point of the mesh (Duprez et al., 2024). The fiber stiffness parameter 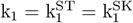 is set equal to 150 kPa following Duprez et al. (2024). In variant ANISO_∥_, anisotropy is related to dermal collagen fibers oriented parallel to the fingertip surface. To achieve this, three local fiber orientations are prescribed at each integration point within the outer layer: the first is aligned with the local meridional direction, and the other two are tilted by +120^°^ and -120^°^ relative to the first. This fiber arrangement ensures transverse isotropy within the outer layer. It also causes traction-compression asymmetry in the membrane as the stiffening obeys the HGO model.
- The most physiologically sound representation of the fingertip tissues is labeled PHYSIO in the next section. It accounts for both the tissue layered heterogeneity and the anisotropy bue to the presence of collagen.

## 4. Results

To establish a common reference for parameter calibration, each model variant is fitted to the mean experimental load–displacement curve up to 1 N contact force. The fit is performed using a custom bounded Hooke–Jeeves (HJ) optimization implemented in Python, minimizing the sum of squared errors between the predicted and experimental load-displacement curves. During calibration, only the shear moduli of the subcutaneous tissues *µ*^ST^ are adjusted. All parameters fall within the typical range used for soft tissue modeling. They are reported in Table 1. The dimensionless collagen stiffening rate parameter k_2_ is fixed to 15 as in Duprez et al. (2024).

**Table 1:**
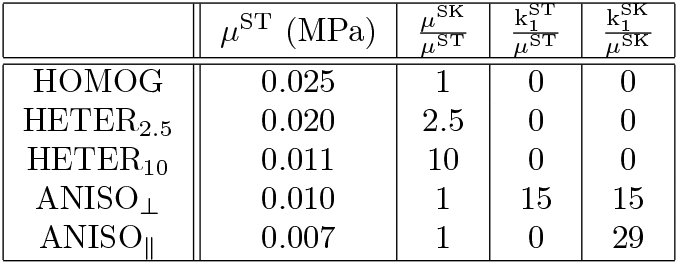
Calibrated material parameters of the HGO model. Shear modulus of the subcutaneous tissues *µ*^ST^; Shear modulus of the skin *µ*^SK^; Stiffness of collagen fibers within the subcutaneous tissues 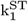 and the skin 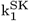.

| | $\mu^{\text{ST}}$ (MPa) | $\frac{\mu^{\text{SK}}}{\mu^{\text{ST}}}$ | $\frac{k_1^{\text{ST}}}{\mu^{\text{ST}}}$ | $\frac{k_1^{\text{SK}}}{\mu^{\text{SK}}}$ |
| --- | --- | --- | --- | --- |
| HOMOG | 0.025 | 1 | 0 | 0 |
| HETER<sub>2.5</sub> | 0.020 | 2.5 | 0 | 0 |
| HETER<sub>10</sub> | 0.011 | 10 | 0 | 0 |
| ANISO<sub>⊥</sub> | 0.010 | 1 | 15 | 15 |
| ANISO<sub>∥</sub> | 0.007 | 1 | 0 | 29 |

Figure 6 (a) presents the force–displacement responses of the different variants. This curve alone does not allow discrimination between the variants, as independent set of parameters can be found for each model to achieve a satisfactory match. Nevertheless, two trends can be identified. On the one hand, isotropic variants display a stiff approximation of the load-displacement response (see HOMOG and HETER_2.5_ and HETER_10_). On the other hand, the introduction of fibers leads to a behavior closer to experimental observations because of the fibers strain-stiffening (see ANISO_⊥_ and ANISO_∥_).

**Figure 6.**
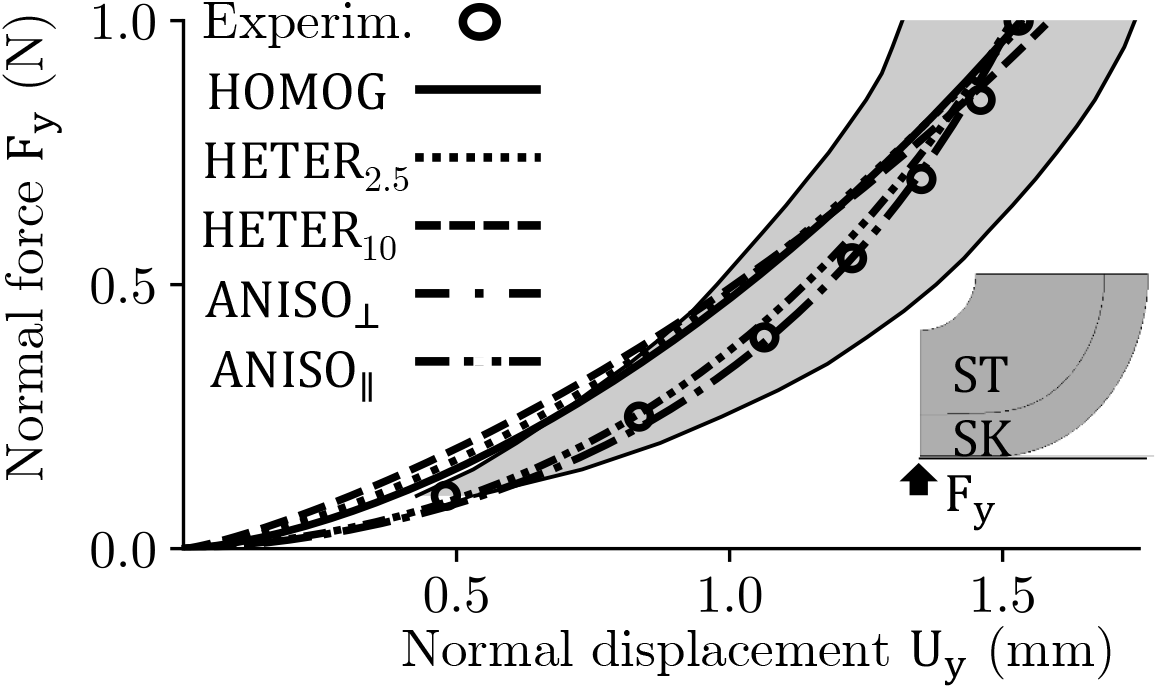
Force–displacement responses. (a) Comparison of the different model variants with the reference experimental curve (with shaded standard deviation); (b) Corresponding calibrated HGO material parameters.

In this Results section, the sensitivity of the model predictions to heterogeneity and anisotropy is investigated. The accuracy of each variant’s predictions is assessed by comparing them with the experimental measurements for two observable quantities. The first quantity corresponds to the surface deformations. Here, the stretches and contractions predicted by each model variant along the meridional and circumferential directions are compared to the principal stretches reported in the experiment. The second quantity is the evolution of the fingertip global shape, as fitted with a spheroid during plate normal loading. The extraction of this measure from the simulations follows the same procedure as for the experimental data described in the previous section (see Figure 2). Subsequently, a comprehensive model is introduced, combining the previously analyzed biomechanical features.

### 4.1. Deformations at the fingertip surface

#### 4.1.1. Influence of heterogeneity

Figure 7 shows the influence of considering that the skin is stiffer than the subcutaneous tissues on the predicted surface stretches by comparing the homogeneous isotropic variant HOMOG with the heterogeneous isotropic variants HETER_2.5_ and HETER_10_. The variation of the local meridional *λ*_m_ and circumferential stretches with increasing meridional angle Ψ is evaluated when the normal force F_y_ = 1 N.

**Figure 7.**
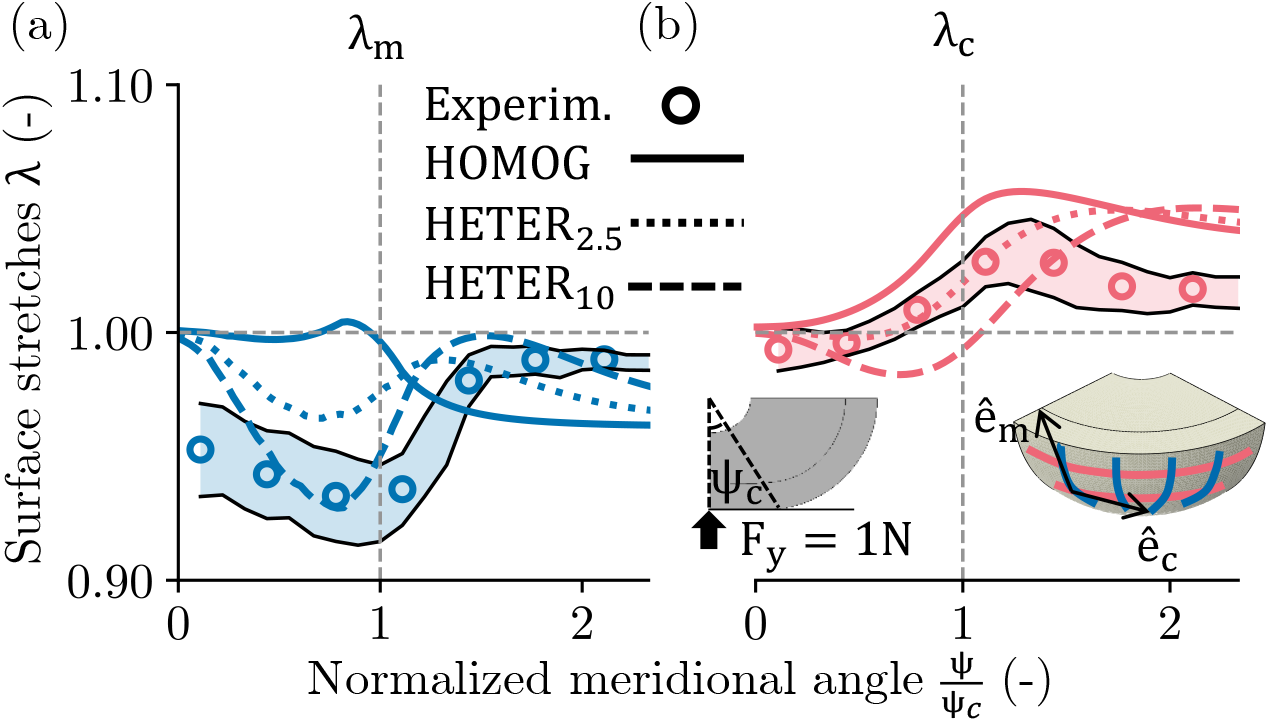
Predictions of the heterogeneity of surface strains under normal load of F_y_ = 1N. (a) Meridional *λ*_m_ and (b) circumferential *λ*_c_ stretches as a function of the meridional angle Ψ; The isotropic homogeneous variant (HOMOG) is compared to the isotropic heterogeneous model variants (HETER) for *µ*^SK^ = 2.5*µ*^ST^ and *µ*^SK^ = 10*µ*^ST^, and to experimental measurements (with shaded standard deviation).

The homogeneous variant displays a realistic reproduction of the circumferential stretches, but does not match the experimental measurements in the meridional direction. This mismatch is particularly striking inside the contact region (i.e., Ψ*/*Ψ_c_ *≤* 1).

Introducing heterogeneity (see HETER_2.5_) slightly improves the accuracy of the predicted meridional stretches both inside and outside the contact region. It also leads to the buildup of meridional stretches near the contact edge (i.e., Ψ*/*Ψ_c_ = 1). The predictions are further improved when the relative stiffness of the outer layer is increased (see HETER_10_). Along the circumferential direction, heterogeneity reduces the magnitude of stretches inside the contact region, which is another improvement over the homogeneous case. However, outside of the contact (i.e., Ψ*/*Ψ_c_ > 1), heterogeneity shifts the maximum circumferential stretch away from the contact edge. Both effects are strengthened when the shear moduli ratio increases (see HETER_10_).

At the contact center (i.e., Ψ*/*Ψ_c_ = 0), all variants predict zero deformation due to the infinite friction coefficient assumption. This is inconsistent with the experimental observations, which show contraction along the meridional direction (see Discussion). Overall, the variant HETER_10_ best reproduces the experimental measurements, with a total SSE of 0.09, compared to 0.12 for HETER_2.5_ and 0.25 for HOMOG.

#### 4.1.2. Influence of subcutaneous and dermal anisotropy

Figure 8 highlights the influence of collagen fiber orientation on the predicted surface stretches. It compares the results of the ANISO_⊥_ model, in which the fibers are oriented radially, with those of the ANISO_∥_ variant, where the reinforcement consists of fibers aligned parallel to the fingertip surface but confined to the outer region of the sphere.

**Figure 8.**
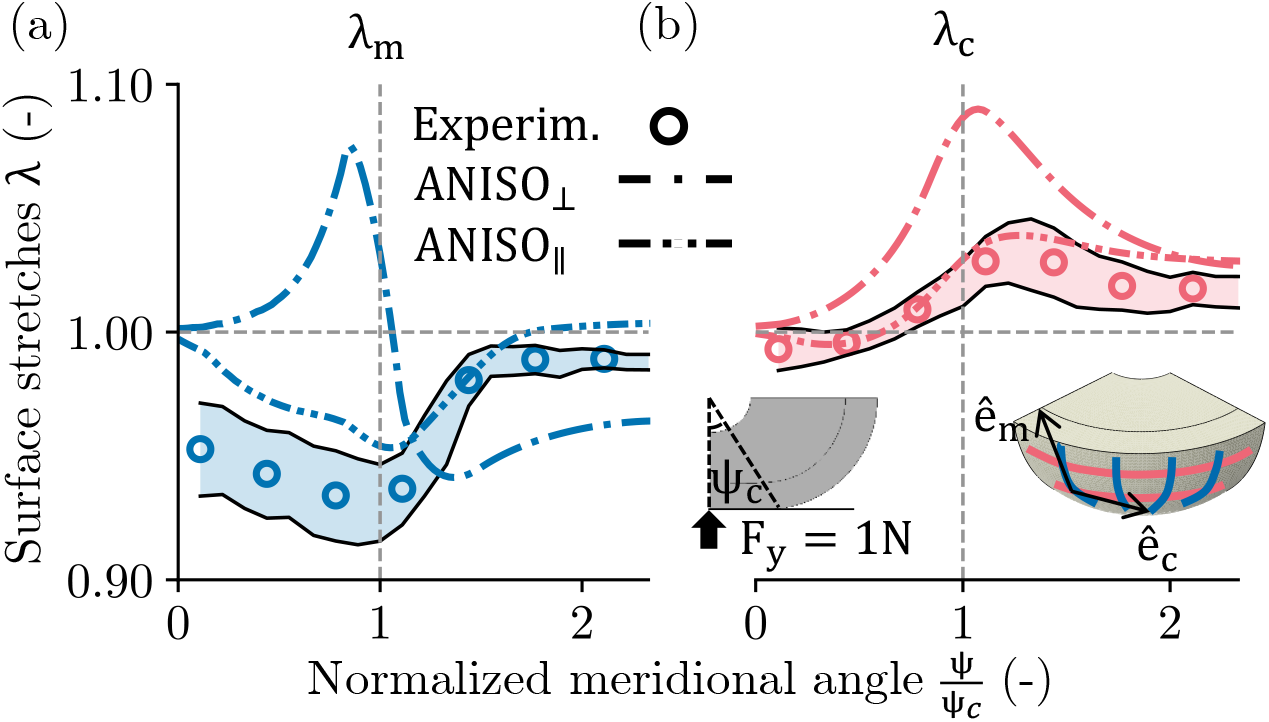
Predictions of the heterogeneity of surface strains under normal load of F_y_ = 1N. (a) Meridional *λ*_m_ and (b) circumferential *λ*_c_ stretches as a function of the meridional angle Ψ; The variant reinforced with radial fibers (ANISO_⊥_) is compared to the variant where the outer layer is reinforced by through-thickness in-plane fibers (ANISO_∥_), and to experimental measurements (with shaded standard deviation).

The predictions of ANISO_⊥_ show similarities with the homogeneous variant (see HOMOG in Figure 7). However, the introduction of radial anisotropy brings the maximum stretches nearby the contact edge (i.e., Ψ*/*Ψ_c_ = 1). Nevertheless, the maximum meridional deformation corresponds to expansion rather than contraction, which does not conform to the experimental measurements.

The introduction of fibers in the outer layer (see ANISO_∥_) leads to realistic predictions, similar to those of the heterogeneous variant (see HETER_10_ in Figure 7). Along the meridional direction, ANISO_∥_ predicts contraction within the contact region (i.e., Ψ*/*Ψ_c_ < 1), with a maximum contraction reached at the contact border (i.e., Ψ*/*Ψ_c_ = 1). The magnitude of the unload outside of the contact (i.e., Ψ*/*Ψ_c_ > 1) is similar to the experimental measurements. Along the circumferential direction, and unlike HETER_10_, reinforcing the outer region with fibers does not move the maximum circumferential stretches away from the contact edge (i.e., Ψ*/*Ψ_c_ = 1). Nevertheless, the heterogeneity of the circumferential stretches is reduced. Overall, the variant ANISO_∥_ best reproduces the experimental data with a total SSE of 0.055 compared to 0.518 for ANISO_⊥_.

### 4.2. Evolution of the pulp overall shape

Figure 9 presents the evolution of the fingertip lateral expansion derived from the spheroid fits. Considering the homogeneous variants HOMOG as baseline, adding a stiff outer layer leads to an increase of the spheroid fit radius during load (see HETER_2.5_ and HETER_10_). On the contrary, introducing radial fibers instead of heterogeneity hinders the growth of the sphere-fit radius (see ANISO_⊥_). Such behavior was observed in our previous study (Duprez et al., 2024). Interestingly, a similar effect is obtained when the outer region is reinforced by fibers parallel to the outer surface (see ANISO_∥_). In this case, however, the reduction in spheroid radius arises from the combined effects of volume preservation and unrealistic surface deformations away from the contact region (see *λ*_m_ in Figure 8), which lead to excessive tissue contraction. The SSE values are 5 for ANISO_∥_, 12 for ANISO_⊥_, 29 for HOMOG, 71 for HETER_2.5_ and 172 for HETER_10_.

**Figure 9.**
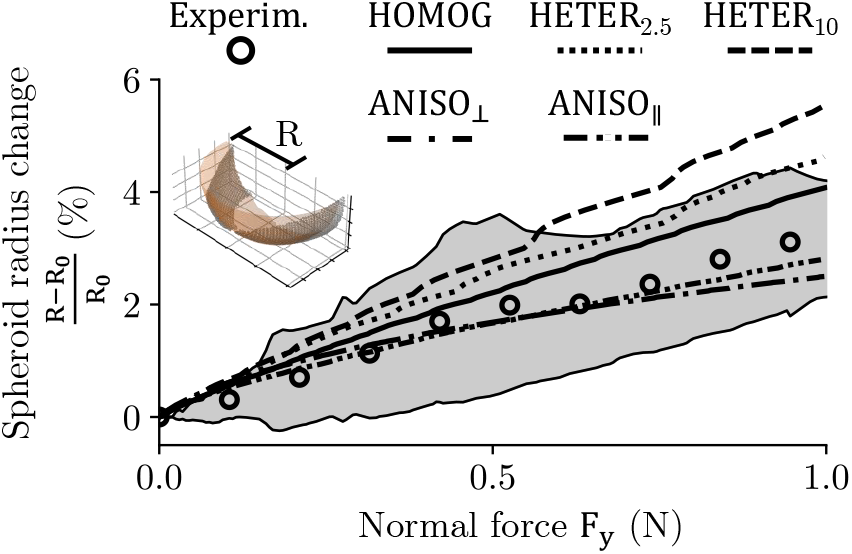
Comparison of the evolution of the fitted spheroid radius derived from experimental 3D point cloud data (with shaded standard deviation) and from numerical simulations.

### 4.3. Subcutaneous deformations

Figure 10 presents the radial *λ*_r_ and meridional *λ*_m_ stretches predicted beneath the fingertip surface by the four model variants. In the subcutaneous tissues, all variants display significant radial contraction (i.e., *λ*_r_ < 1) below the contact region and radial expansion (i.e., *λ*_r_ > 1) elsewhere. Conversely, along the meridional direction, subcutaneous tissues expand below the contact (i.e., lambda_m_ > 1) and contract in the surrounding regions (i.e., *λ*_m_ < 1). The inclusion of radial fibers reduces subcutaneous expansion (see ANISO_⊥_).

**Figure 10.**
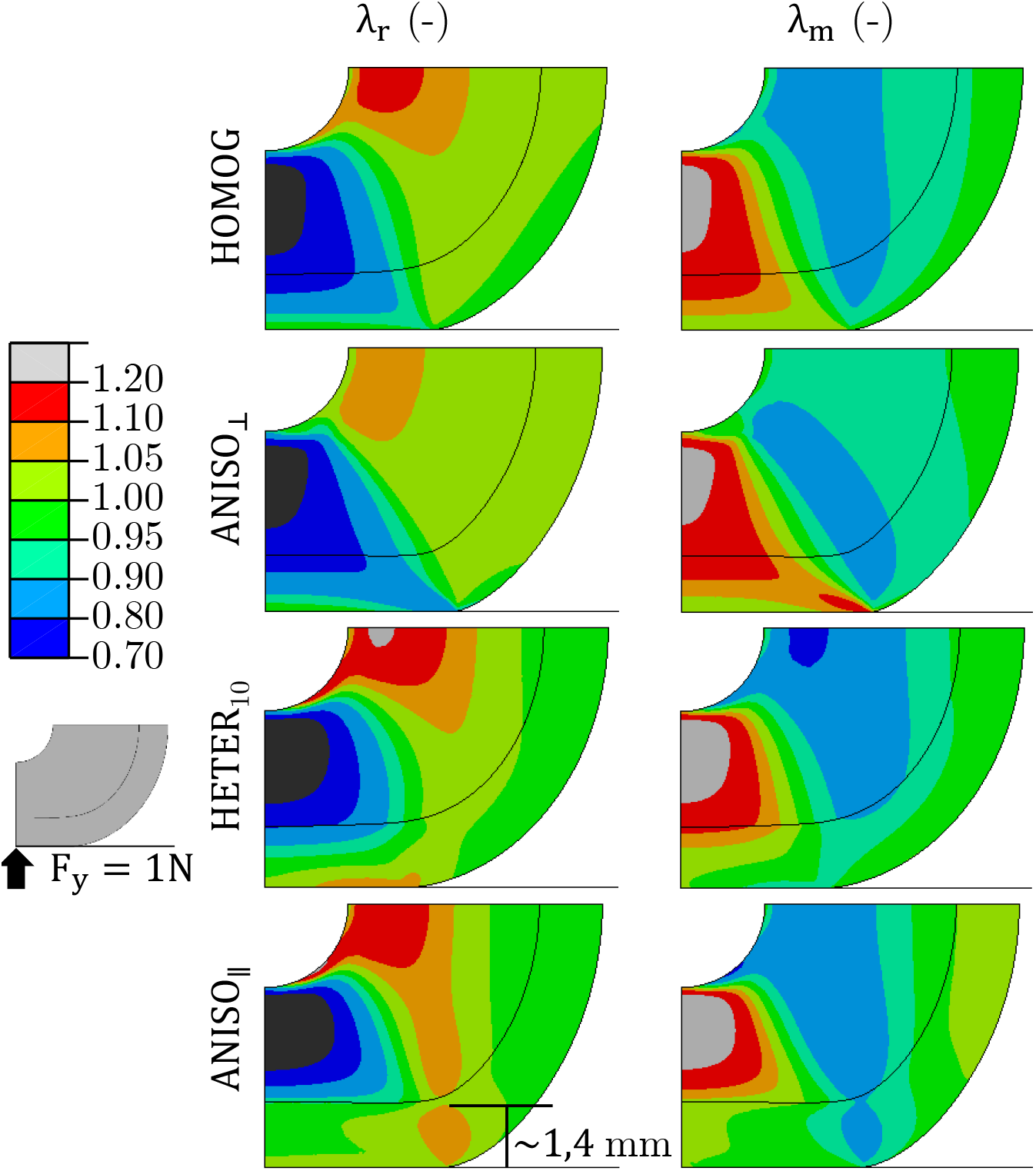
Subsurface field of the local radial *λ*_r_ and meridional *λ*_m_ stretches predicted by the different variants of the model under a normal load of F_y_ = 1N.

In the skin, variations in local thickness reflect the degree of material heterogeneity. Beneath the contact region, the homogeneous cases (i.e., HOMOG and ANISO_⊥_) exhibit thinning (i.e., *λ*_r_ < 1), whereas thickening occurs when heterogeneity is introduced (i.e., *λ*_r_ > 1, see HETER_10_). Along the meridional direction, heterogeneity causes tissue contraction beneath the contact region. A comparable trend is observed when heterogeneity is modeled via the inclusion of in-plane fibers in the outer layer rather than by introducing a contrast in shear moduli between layers (see ANISO_∥_). Away from the contact, these heterogeneous variants predict a reduction in skin thickness. Finally, only ANISO_∥_ predicts significant and deep tissue thickening (i.e., *λ*_r_ > 1) beneath the contact edge. This variant also predicts meridional expansion below the surface (i.e., *λ*_m_ > 1) away from the contact region.

### 4.4. Towards a realistic representation of the tissues

Figure 11 presents the comprehensive fingertip model, labeled PHYSIO. It incorporates a stiff outer skin layer reinforced by a network of collagen fibers oriented parallel to the outer surface and with a subcutaneous network of radial collagen fibers. As for the previous variants, the material parameters are calibrated to load-displacement data using HJ optimization, and it is now done in two steps. Starting from the calibrated parameters of the variant ANISO_∥_, the fibers stiffness parameter in the outer layer 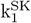 is decreased from 200 kPa to 150 kPa while the shear modulus of the isotropic background matrix in the outer layer *µ*^SK^ is adjusted to maintain the load-displacement behavior. Similarly, the shear modulus of the inner layer *µ*^ST^ is then halved, and the subcutaneous fibers stiffness parameter 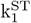 is adjusted. The predicted load-displacement response is shown in Figure 11 while the corresponding calibrated material parameters are *µ*^ST^ = 0.004 MPa and 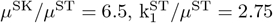 and 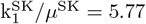. All parameters fall within the typical range used for soft tissue modeling.

**Figure 11.**
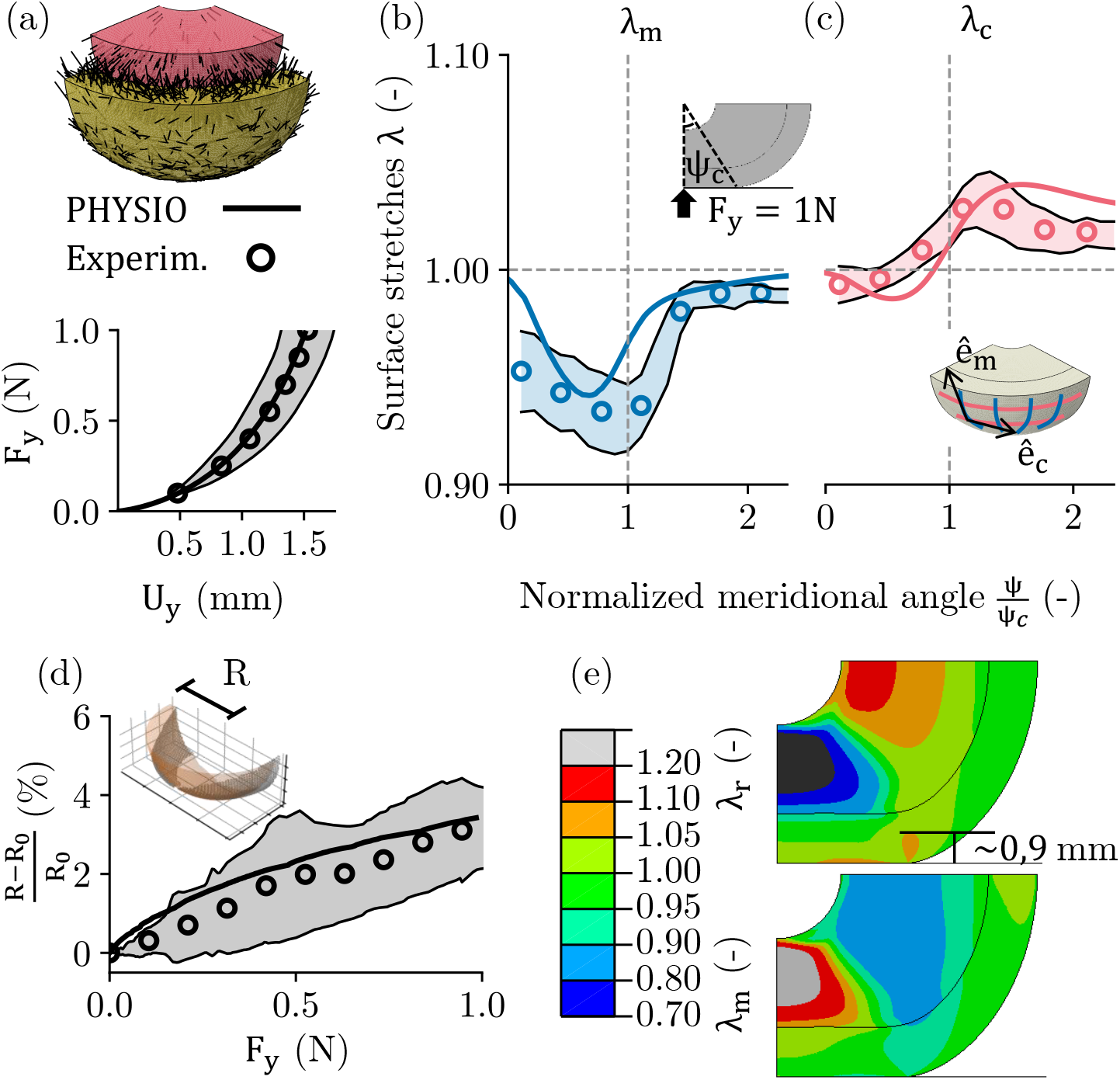
Assessment of the comprehensive model compared to experimental measurements (with shaded standard deviation) under a normal load of F_y_ = 1 N. (a) Force-displacement response; (b) Meridional and (c) circumferential surface stretches; (d) Evolution of the fitted spheroid radius; (e) Subcutaneous radial and meridional deformations.

Figure 11 (b) and (c) show the surface meridional and circumferential stretches predicted by the physiological variant. Accurate reproduction of the surface deformations are obtained along both directions, with localization of strains around the contact border (i.e., Ψ*/*Ψ_c_ = 1). A realistic prediction of the evolution of the pulp overall shape is also obtained (see Figure 11 (d)). Below the contact (see Figure 11 (e)), the variant predicts a gradient of radial stretch with expansion (i.e., *λ*_r_ > 1) just below the surface, and with a transition to contraction (i.e., *λ*_r_ < 1) roughly halfway through the skin layer. A sharp radial expansion still arises close to the contact edge and propagates 0.9 mm deep. The SSE values for the comprehensive model are 0.052 for the surface strain predictions and 24.38 for the global shape predictions.

## 5. Discussion

The predictions of the different model variants highlight the significant role of tissue heterogeneity, anisotropy, and traction-compression asymmetry in shaping fingertip deformation during normal loading. As expected, considering a stiffer outer layer (i.e., skin) improves the agreement with measured surface strains (see Figure 7 and 8). However, more realistic behaviors are only obtained with a large contrast. This suggests that the widely used stiffness ratio of 2.5 (Serhat et al., 2022; Talarico et al., 2014; Lesniak and Gerling, 2009) underestimates the ability of the skin to constrain the pulp. The 2.5 ratio originates from small-scale indentation tests (200 *µ*m depth, 100 *µ*m indenter) (Maeno et al., 1998), which mainly probe the background matrix rather than engaging the collagen network (Wahlsten et al., 2023). As loading increases, collagen fibers progressively align with the applied deformation, producing strong strain stiffening (Haut, 2002). Hence, for large-scale deformations, modeling heterogeneity solely through a 2.5 stiffness ratio is likely unrealistic.

The contrast in material properties is also shown to induce a thickening of the skin below the contact region (see Figure 10). The thickening of the stratum corneum (i.e., the outermost skin layer) during normal loading has been observed experimentally using OCT imaging of the fingerpad (Corniani et al., 2025). Models treating skin as a membrane (Srinivasan, 1989) cannot capture such local thickness variation. Still, membrane models can approximate fingertip surface deformations during flat plate contact (Kumar and DasGupta, 2013). However, such models predict a more diffuse meridional stretch distribution and fail to reproduce the circumferential stretch maximum at the contact edge (Doumont et al., 2025).

While heterogeneity influences the localization of strains around the contact edge (see Figure 7), our results show that anisotropy and traction-compression asymmetry are also important (see Figure 8). Because collagen fibers carry virtually no compressive load compared to tensile load, it has been shown that radial fibers flatten the contact pressure profile (Duprez et al., 2024), consistent with experimental observations (Johansson and Flanagan, 2009). This redistribution of load towards the contact edge is reflected in sharper surface deformation gradients at the transition between contact and free surface.

Given the relative magnitudes of the experimental meridional and circumferential stretches at the contact edge, local skin thickening is expected as a consequence of volume preservation. This is predicted when considering heterogeneity, but deeper skin stretches occur when introducing dermal collagen fibers (see Figure 10). In the case of the physiological model (see Figure 11 (e)), this thickening reaches the SAI and RAI mechanoreceptors which are reported at *∼*1 mm below the surface (Deflorio et al., 2022). This suggests dermal fibers may facilitate the propagation of surface strains into the vicinity of sensory units. The role of collagen in the deep transmission of mechanical stimuli has already been proposed (Tanaka et al., 2015). Our results provide further support for this hypothesis.

Despite the comprehensive model accurately reproducing the fingertip behavior during normal loading, several modeling simplifications were adopted in this study. The first is the hypothesis of axisymmetry. While it is reasonable at the macroscopic scale, this assumption no longer holds at finer scales. At the microscale, the fingertip is not perfectly smooth. Its surface ridges have been shown to induce frictional anisotropy between the proximal–distal and radial–ulnar directions through an interlocking mechanism (Chimata and Schwartz, 2015).

The frictional behavior is also simplified by assuming no relative motion between contacting surfaces once contact is established (i.e., *µ* =*∞*). However, the experimental results show meridional contraction of the skin surface around the contact center, suggesting that partial slip occurs within the contact region (see Experim. in Figure 7) (Doumont et al., 2025). The fingerprint ridges, which induce frictional anisotropy through interlocking mechanism (Chimata and Schwartz, 2015), have also been neglected. While such effects may affect the shape of the contact area, their impact during normal loading is expected to be negligible. Incorporating a simple Coulomb friction law should improve the reproduction of experimental measurements.

The representation of collagen fibers within the skin has also been simplified. Through-thickness fibers (Hauck et al., 2004) were omitted, although this is reasonable given that dermal collagen is predominantly oriented in-plane (Munisso et al., 2023). Moreover, no preferential in-plane orientations were assigned, despite evidence of two dominant fiber directions in some skin sites (Munisso et al., 2023; Silver, 2022). However, most of these observations were not made on fingertip specimens, and collagen orientation is known to be site-specific (Ní Annaidh et al., 2012). Therefore, modeling the skin as transversely isotropic with asymmetric traction–compression behavior in-plane remains a reasonable and realistic assumption.

Finally, the nail has not been considered, although some mechanoreceptor endings lie in its neighborhood (Birznieks et al., 2009). Rather, the vertical displacement of the top surface has been impeded. Nevertheless, previous fingertip models including the nail showed little displacement in its vicinity (Talarico et al., 2014; Wu et al., 2006). Moreover, it was fixed in the reference experiment (Doumont et al., 2025). Consequently, the boundary condition on the top surface appears reasonable.

## 6. Conclusions

To investigate the influence of soft tissue biomechanics on fingertip surface strains, we developed a 3D FE model of fingertip contact with a glass plate under normal loading. The predictions conform to experimental measurements of fingertip surface strains, lateral expansion, and load-displacement response. A sensitivity analysis showed the respective roles of tissue heterogeneity and collagen fibers. Accurate predictions were achieved when both the contrasted properties of the skin and subcutaneous tissue, and the anisotropy due to collagen fibers were taken into account:

- In the subcutaneous tissue, radial collagen fibers prevented unrealistic fingertip expansion during loading and enabled the localization of circumferential strains near the contact edge, consistent with experimental observations;
- In the skin, collagen fibers oriented parallel to the fingertip surface provided in-plane stiffness, limiting excessive circumferential elongation and promoting skin thickening, as also reported in experimental measurements.

## Declaration of generative AI and AI-assisted technologies in the manuscript preparation process

During the preparation of this work, the authors used Microsoft Copilot and OpenAI’s ChatGPT to assist in the design of Python code for optimization schemes and figure generation from simulation data. After using these tools, the authors reviewed and edited the content as needed and take full responsibility for the content of the published article.

### Sample CRediT author statement

**Guillaume H. C. Duprez:** writing – original draft, review & editing, conceptualisation, methodology, software, visualisation **Donatien Doumont:** writing – review & editing, resources, visualisation **Philippe Lefèvre:** writing – review & e-diting **Benoit P. Delhaye:** writing – review & -editing, conceptualisation, methodology, supervision **Laurent Delannay:** writing – review &- editing, conceptualisation, methodology, supervision, funding acquisition.

## Acknowledgments

Computational resources have been provided by the Consortium des Équipements de Calcul Intensif (CÉCI), funded by the Fonds de la Recherche Scientifique de Belgique (F.R.S.-FNRS) under Grant No. 2.5020.11 and by the Walloon Region. LD, BD and PL are mandated by the F.R.S.-FNRS.

